# Interval Timing Shows Selective Enhancement Under Psychosocial Stress: A Cortisol-Mediated Dissociation From Spatial Processing

**DOI:** 10.1101/2025.11.07.686532

**Authors:** Güvem Gümüş-Akay, Gözde Vatansever, Filiz Çetinkaya, Nuray Varol, Metehan Çiçek

## Abstract

Acute stress affects cognitive processes, including our perception of time, yet few studies have examined how HPA axis activation specifically modulates time perception. This study provides the first systematic examination of how psychosocial stress influences temporal versus spatial reproduction using the Trier Social Stress Test (TSST), investigating underlying epigenetic mechanisms. Forty-four healthy adults (21 men; mean age 21.96 ± 3.03) completed temporal and spatial reproduction tasks before and after TSST, with salivary cortisol sampling and DNA methylation analysis of four dopamine-related genes (*COMT, DRD2, SLC6A3, TH*).

The TSST successfully elevated cortisol and state anxiety, confirming effective HPA axis activation. Critically, stress selectively reduced temporal underestimation. Despite their shared neurocognitive mechanisms, spatial processing remained unchanged. This demonstrates that HPA-mediated stress enhances interval timing. Men demonstrated greater stress-induced improvement in temporal accuracy than women, while neither sex showed significant spatial changes, independent of cortisol reactivity differences. The results are discussed in the context of dopaminergic models of temporal processing.

Exploratory epigenetic analyses revealed a *DRD2* methylation × cortisol response interaction for temporal tasks, with higher methylation associated with greater stress-induced improvement among high cortisol responders. However, sensitivity analysis indicated these interactions were driven by participants with extreme methylation values, limiting generalizability of epigenetic results.

These findings demonstrate that acute psychosocial stress selectively enhances temporal accuracy, potentially through cortisol-dopamine interactions in corticostriatal timing circuits. This work opens avenues for investigating stress-timing mechanisms and cortisol-dopamine interactions in timing circuits, and provides preliminary evidence for epigenetic moderation.

## 1. Introduction

The sense of time is tracked using a variety of mechanisms across species, with timescales ranging from milliseconds to days (Buhusi and Meck, 2005). The human brain can monitor time continuously and guide complex motor actions that require millisecond precision (Lewis and Walsh, 2005; Merchant et al., 2013). “Interval timing” refers to durations in the seconds-to-minutes range, but can also describe shorter (millisecond) and longer (hour) intervals (Meck et al., 2003). A range of basic cognitive and emotional factors including attention, sensory modality, arousal and emotional load systematically influences interval timing (Droit-Volet and Meck, 2007). These influences largely depend on the underlying neural mechanisms that support time perception.

Interval timing relies heavily on dopaminergic signalling. Converging lesion and neuroimaging evidence implicates basal ganglia circuits in interval timing (Coull et al., 2011). Consistent with this, robust impairments in timing have been shown by disorders marked by dopamine dysfunction, such as Parkinson’s disease (Merchant et al., 2008; Smith et al., 2007). Furthermore, pharmacological manipulation of the dopaminergic system has been found to reliably shift perceived duration (Rammsayer, 1989; Rammsayer, 1993, 1997; Rammsayer and Vogel, 1992). Lastly, optogenetic manipulations provide causal evidence: selective modulation of substantia nigra pars compacta dopamine neurons biases interval judgements toward over- or underestimation (Soares et al., 2016). Although these circuits specify how timing is processed, the system’s estimates can be systematically biased by internal states and environmental threats.

Recent research demonstrates that negative or aversive stimuli presented across different sensory modalities can disrupt interval timing. A well-documented effect is that time often appears to pass more slowly than it actually does when individuals are confronted with threatening, unpleasant, or fear-inducing events, a phenomenon commonly referred to as time dilation (Bar-Haim et al., 2010; Droit-Volet et al., 2004). This distortion of temporal experience has been linked to emotion-driven psychophysiological changes (Craig, 2009), suggesting that stressful experiences which trigger physiological arousal may alter time perception. Consistent with this view, it was reported that elevated skin conductance responses (reflecting the activation of the sympathetic-adrenal-medullary system; SAM) were associated with time dilation in response to negative stimuli (Mella et al., 2011). Van Hedger et al. (2017) examined the relationship between psychophysiological stress responses and interval timing. They observed that social stress induced time dilation; however, the observed distortion in time perception was not fully explained by the autonomic nervous system reactivity, and thus SAM-axis activity (van Hedger et al., 2017).

This brings into focus the second major stress-response pathway, the hypothalamic-pituitary-adrenal (HPA) axis. Unlike the fast-acting SAM system, the HPA axis unfolds more slowly (Wirth, 2015; Craig, 2009), resulting in the release of glucocorticoids which regulate energy redistribution and influence neural activity. Importantly, these signalling cascades are not only transient physiological adjustments. Cortisol crosses the blood-brain barrier and, given the widespread distribution of glucocorticoid receptors (GRs) in the brain (Nelson and Kriegsfeld, 2017), it can affect various brain functions; furthermore, it can also bind intracellularly, translocate to the nucleus, and regulate transcriptional programs. Taking these broad effects into account, cortisol emerges as the key candidate molecule in the mediation of the cognitive and behavioural effects of stress. Meanwhile, stress-related pathways intersect with dopaminergic signalling in corticostriatal circuits that support temporal judgement and cognitive control. While such intersections between glucocorticoid and dopaminergic signalling may highlight a plausible mechanism for stress-related distortions of time, they raise the further question of what accounts for individual differences in these outcomes.

Individuals differ substantially in their perception of time, just as they do in their stress reactivity. Prior genetic research indicates that variation at the DNA sequence level alone cannot fully account for this diversity (Bartholomew et al., 2015; Sysoeva et al., 2010). This suggests that epigenetic mechanisms, which are shaped by the interaction between genes and the environment, may be key contributors. However, to our knowledge, no studies have directly examined how DNA methylation relates to time perception.

Building on this gap, we conducted an experiment that jointly examined stress physiology, temporal cognition, and molecular markers. We employed the Trier Social Stress Test (TSST), a well-established in-lab stress induction method (Allen et al., 2017; Dickerson and Kemeny, 2004) to trigger psychosocial stress. Temporal and spatial reproduction tasks, matched for sensory and motor demands, and neurocognitive resources (Skagerlund et al., 2016; Üstün et al., 2022; Walsh, 2003), were administered before and after TSST. Serial salivary cortisol sampling indexed HPA axis activation and was used to classify participants into high versus low cortisol increase rate (CIR) groups. In parallel, we assessed salivary DNA methylation of four dopamine-related genes with distinct functional roles: *COMT* (dopamine degradation), *DRD2* (dopamine D2 receptor), *SLC6A3*/DAT (dopamine transporter), and *TH* (dopamine synthesis). Behavioral responses were analyzed using a linear mixed-effects framework with stress condition, task type, sex, and CIR group as factors, controlling for age and body mass index (BMI). This multi-level approach enabled us to address three key aims: (i) to test whether acute psychosocial stress alters temporal versus spatial reproduction in a domain-specific manner; (ii) to determine whether cortisol responder status modulates these effects; and (iii) to examine whether methylation of dopamine-related genes moderates stress-induced changes in reproduction performance, and whether such effects are specific to time or extend to space.

## 2. Methods

### 2.1. Participants

Interested applicants completed an online survey to assess eligibility and gather demographic information, including the Turkish versions of the State-Trait Anxiety Inventory-Trait Form (Spielberger, Gorduch and Lushene 1970; Öner and Le Compte 1998) and Handedness Scale (Chapman and Chapman, 1987; Nalçacı et al., 2002). To be eligible, participants had to be between 18 and 35 years old, right-handed, and have no history of neuropsychiatric or endocrine diseases or regular medication use. We excluded individuals who worked night shifts or had recent COVID-19 infection.

Initially, 46 healthy, right-handed volunteers participated. Two participants were unable to finish the TSST, resulting in a sample of 44 participants (21 men; mean age 21.96 ± 3.03 years). Women participants were scheduled during the luteal phase of their menstrual cycle, and none were taking oral contraceptives. Participants followed standard pre-test restrictions (no exercise, caffeine, or food 24h prior) and completed screening questionnaire, proposed for the standardization of the TSST protocol (Narvaez Linares et al., 2020).

All participants provided informed consent. The study was approved by the Ankara University Faculty of Medicine Human Research Ethics Committee (approval # İ3-104-19) and aligned to the principles of the Declaration of Helsinki.

### 2.2. Experimental Procedure

We conducted all sessions between 13:30 and 16:30 to control for diurnal cortisol variations. Before the experiment began, participants were given a few minutes to rest in the behavioural testing room. They were given a brief explanation of the study procedures and saliva sample collection. The session began with the first saliva sample being taken (t_-30_: baseline). Participants completed the state form of the State-Trait Anxiety Inventory (STAI-State: pre-TSST) when they provided the saliva sample. The TSST Background Questionnaire was also administered at this stage.

To assess the general cognitive functions, Forms A and B of the Trail Making Test (TMT) were administered sequentially, with completion times and error scores recorded for each form. As elevated cortisol levels may influence outcomes, the TMT was conducted in the pre-TSST phase. Participants then began the experimental paradigm, completing the computer-based reproduction task which took about 10 minutes (see the next chapter for details of the task). At the end of the initial 30-minute pre-stress phase, a second saliva sample was collected (t_0_). Participants were then escorted to another room to undergo the TSST, which was administered without prior notification of its content.

Fifteen minutes after entering the TSST room, participants returned to the first room, where a third saliva sample (t+15) was collected and they completed the STAI-State for a second time (post-TSST). Between t_+15_ and t_+30_ samples, the time perception paradigm was conducted again, for the post-stress measurement. The remaining psychometric scales were then administered. Additional saliva samples were collected at t_+30_, t_+45_ and t_+60_. Full details of all psychometric instruments, their descriptive statistics and the TSST procedure can be found in the Supplementary Information.

### 2.3. Reproduction Task Paradigm

The task was based on a reproduction method, in which participants first perceived a target stimulus defined by either its duration or size, and then manually reproduced it. The inclusion of a matched spatial perception condition as a control was for the purpose of controlling for general magnitude processing. Each trial began with a fixation cross on a grey background, followed by an instruction cue. An hourglass icon indicated that the upcoming trial required reproduction of stimulus duration, while a ruler icon indicated reproduction of stimulus size. In both conditions, a black circle appeared in the center of the screen. Its radius (100, 125, 150, 175 or 200 pixels) and display duration (2, 2.5, 3, 3.5 or 4 seconds) varied, and the task cue specified which dimension participants had to reproduce.

In the time condition, participants attended to the duration for which the circle remained visible (Figure 1A). After it disappeared, they pressed and held the left mouse button to make the circle reappear, releasing it when they judged that the elapsed time matched the target duration. The time between pressing and releasing the button was recorded as the reproduced time. In the space condition, participants judged the size of the circle. After the target circle disappeared, they pressed and held the mouse button to reveal a fixed, smaller circle whose radius gradually increased while the button remained pressed. They released the button once they estimated that the circle matched the target size. The final radius at button release was recorded as the reproduced size.

**Figure 1.**
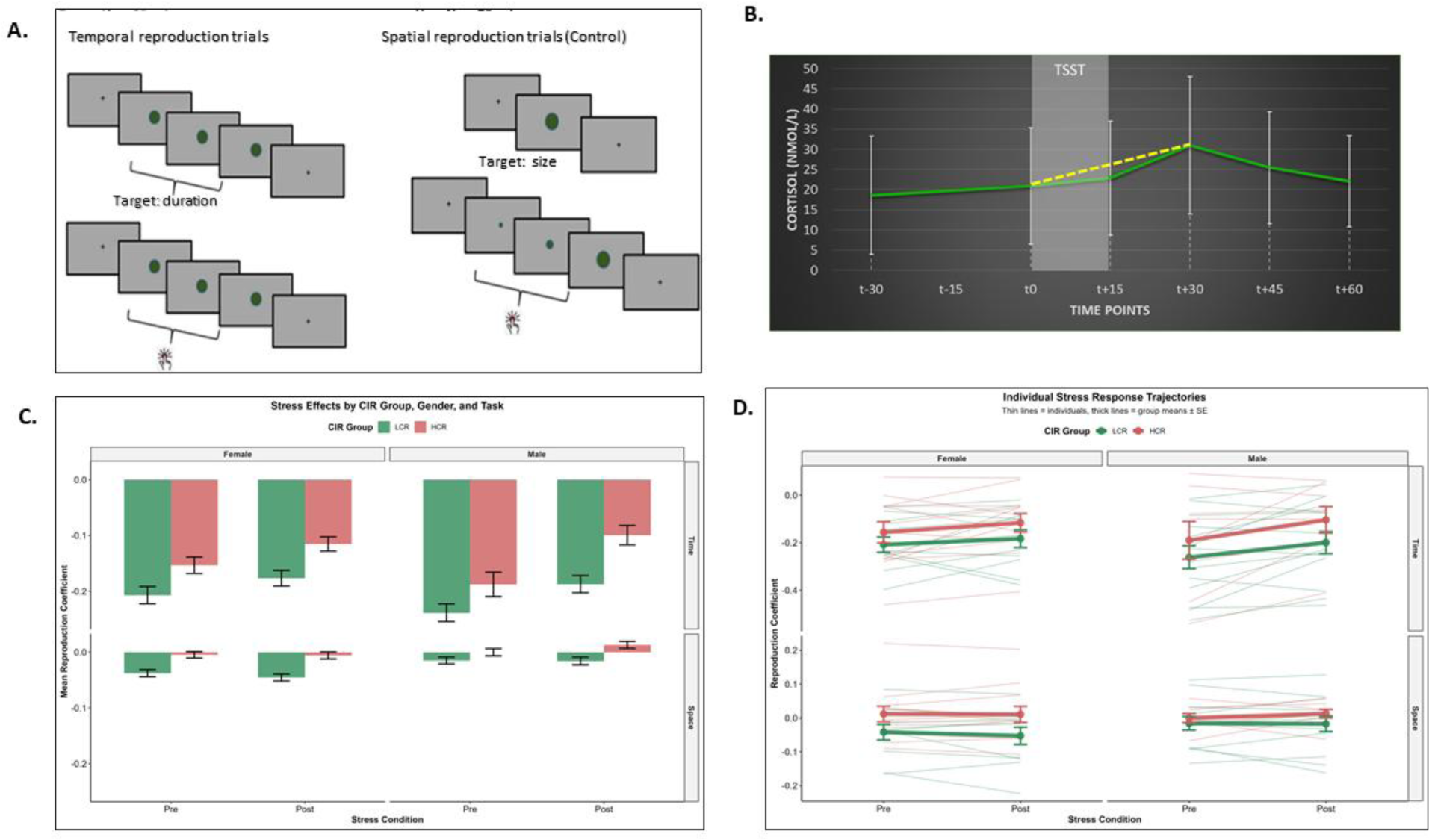
**A.** Schematic representation of the reproduction task with temporal and spatial trials. **B.** Cortisol response curves of participants, derived from five saliva points. First application of the reproduction task (pre-stress) was between t_-30_ and t_0_, TSST was conducted between t_0_ and t_+15_, second reproduction task (post-stress) took place between t_+15_ and t_+30_ samples. Yellow line indicates the cortisol increase rate (CIR). **C.** Reproduction coefficients for each task, condition, group and sex. **D.** Individual trajectories of participants’ behavioral responses before and after TSST. Thick lines show group mean.

The paradigm comprised 50 randomised trials per session (25 temporal and 25 spatial), with each of the five combinations of target duration and target radius presented five times. The task was administered twice, before and after the TSST, to enable assessment of stress-related changes in temporal versus spatial reproduction. The stimuli were presented on a 15.6-inch monitor with a resolution of 1920 x 1080 and a refresh rate of 60 Hz, with participants seated approximately 60 cm from the screen.

Using the same approach as in the other reproduction studies, we computed a coefficient of variation (CV) to enable direct comparison of reproduction performance across time and space tasks, which provided a common metric for both dimensions. The CV quantified both the magnitude and the direction of error, capturing how far the reproduced value deviated from the target value. The CV was calculated for each trial using the following formula: *CV= 1 - (Target Value/Reproduced Value)*(Barkley and Fischer, 2019). A CV of 0 indicated a reproduction of perfect accuracy. Negative CV values indicated shorter or smaller reproductions relative to the target (underestimation), while positive CV values indicated longer or larger reproductions (overestimation). Descriptive statistics can be found in Table 1.

**Table 1.** Descriptive statistics of the behavioral responses (reproduction coefficients).

| Reproduction Coefficient |  |  |  |  |  |
| --- | --- | --- | --- | --- | --- |
| Task Type | Condition | Group | N | Mean (SD) | Median [IQR] |
| Time | Pre | LCR | 547 | -0.224 (0.26) | -0.18 [-0.371--0.046] |
| Time | Pre | HCR | 470 | -0.167 (0.269) | -0.123 [-0.312-0.034] |
| Time | Post | LCR | 560 | -0.182 (0.249) | -0.153 [-0.316--0.015] |
| Time | Post | HCR | 488 | -0.109 (0.229) | -0.102 [-0.223-0.059] |
| Space | Pre | LCR | 579 | -0.025 (0.107) | -0.016 [-0.097-0.049] |
| Space | Pre | HCR | 468 | -0.003 (0.093) | -0.002 [-0.062-0.057] |
| Space | Post | LCR | 577 | -0.029 (0.115) | -0.022 [-0.105-0.05] |
| Space | Post | HCR | 469 | 0.002 (0.097) | 0.006 [-0.064-0.075] |

**Table 2.** Mixed-effect model results.

| Effect | F | Df<sub>1;2</sub> | p | $\eta^2_p$ | f |
| --- | --- | --- | --- | --- | --- |
| Stress | 21.7195 | 1;4,102.44 | <0.0001 | 0.0053 | 0.0728 |
| Group | 4.5008 | 1;37.93 | 0.0405 | 0.1061 | 0.3445 |
| Sex | 0.0056 | 1;37.98 | 0.9406 | 0.0001 | 0.0122 |
| Task | 790.4374 | 1;4,108.31 | <0.0001 | 0.1614 | 0.4386 |
| Age | 0.5031 | 1;37.98 | 0.4825 | 0.0131 | 0.1151 |
| BMI | 0.1333 | 1;37.97 | 0.7170 | 0.0035 | 0.0593 |
| Stress:Group | 2.2538 | 1;4,102.46 | 0.1334 | 0.0005 | 0.0234 |
| Stress:Sex | 4.4559 | 1;4,102.40 | 0.0348 | 0.0011 | 0.0330 |
| Group:Sex | 0.0015 | 1;38.01 | 0.9694 | 0.0000 | 0.0063 |
| Stress:Task | 21.7546 | 1;4,102.32 | <0.0001 | 0.0053 | 0.0728 |
| Group:Task | 10.7309 | 1;4,108.45 | 0.0011 | 0.0026 | 0.0511 |
| Sex:Task | 12.2975 | 1;4,108.41 | 0.0005 | 0.0030 | 0.0547 |
| Stress:Group:Sex | 0.4097 | 1;4,102.43 | 0.5222 | 0.0001 | 0.0100 |
| Stress:Group:Task | 0.2201 | 1;4,102.36 | 0.6390 | 0.0001 | 0.0073 |
| Stress:Sex:Task | 1.1711 | 1;4,102.37 | 0.2792 | 0.0003 | 0.0169 |
| CIR_Group:Sex:Task | 2.1067 | 1;4,108.29 | 0.1467 | 0.0005 | 0.0226 |
| Stress:Group:Sex:Task | 0.2500 | 1;4,102.37 | 0.6171 | 0.0001 | 0.0078 |

### 2.4. Trier Social Stress Test and Cortisol Response

To activate the HPA axis, we used TSST, an experimental setup with high ecological validity for investigating psychobiological stress responses in a laboratory setting (Allen et al., 2017; Birkett, 2011). Briefly, participants are required to deliver an unprepared speech in front of an unresponsive jury, which elevates cortisol through key stress factors such as social evaluation, unexpected events (e.g., a surprise arithmetic question) and public speaking.

Saliva cortisol levels were determined by the competitive ELISA method using a Cortisol Saliva ELISA kit (DiaMetra, DKO020) according to the manufacturer’s instructions. Each sample was analysed in duplicate. Cortisol response curves for each participant were generated by plotting cortisol concentrations against six different time points. To convert the participants’ cortisol responses obtained at different time points into single quantitative data, cortisol increase rate (CIR) values were calculated for the t_0_–t_+30_ time interval using the following formula: formula: CIR = [Cortisol]t_+30_-[Cortisol]t_0_/time (Khoury et al., 2015). Participants were divided into two groups: *low cortisol responders* (LCR, n = 24) and *high cortisol responders* (HCR, n = 20). Participants were assigned to these groups based on their CIR values, using the overall group mean value (0.34) as the cut-off point.

### 2.5. DNA methylation analysis

Genomic DNA was extracted from saliva using the MasterPure Complete DNA and RNA Purification Kit (Lucigen, MC85200) according to the manufacturer’s protocol. Bisulfite modification of 500 ng DNA samples was performed using the EZ-DNA Methylation Gold Kit (Zymo Research, D5006) according to the manufacturer’s instructions. The regulatory regions of target genes were amplified using methylation-specific primers (Supplementary Table 1), with one primer biotin-labeled (Qiagen, 978746), and the PyroMark PCR kit (Qiagen, 978703). PCR cycling conditions consisted of initial denaturation at 95 °C for 15 min, followed by 45 cycles of 95 °C for 30 s, 60 °C for 30 s, and 72 °C for 30 s, with a final extension at 72 °C for 10 min. Each experiment included bisulfite-modified methylated and unmethylated human DNA controls (Qiagen, 59695) and no-template controls. PCR products were verified by 2% agarose gel electrophoresis before sequencing. Methylation profiles of CpG loci were determined by pyrosequencing using the PyroMark Q24 System (Qiagen, Germany) with sequencing primers listed in Supplementary Table 1, following the manufacturer’s workflow. Methylation levels (% methylation) were analyzed using PyroMark Q24 2.0.8 software. Descriptives for methylation levels are provided in Supplementary Table 4.

### 2.6. Statistical analysis

All analyses were conducted using R Studio (2025.05.1; R version 4.4.1) with the following packages: tidyverse, lmerTest, emmeans, and effectsize. Statistical significance was set at α = 0.05, and effect sizes were interpreted according to Cohen’s conventions (small: f ≥ 0.1, medium: f ≥ 0.25, large: f ≥ 0.4). Reproduction coefficients were calculated using the formula stated above for both temporal and spatial dimensions. Outliers were identified and removed using the interquartile range method (values beyond 1.5 × IQR from Q1 and Q3) separately for each task type. Participants were categorized into high cortisol increase rate (HCR) and low cortisol increase rate (LCR) groups based on their stress response profiles.

The main statistical model employed a linear mixed-effects approach using the lmer function from the lmerTest package. This 2×2×2×2 factorial design examined the four-way interaction between stress condition (Pre vs. Post), CIR group (LCR vs. HCR), sex (Women vs. Men), and task type (Time vs. Space), while controlling for age and BMI as covariates. Model assumptions were verified through visual inspection of residual plots. Residuals showed approximately normal distribution and homogeneous variance across fitted values. Partial eta-squared (η²p) was calculated for all effects. Cohen’s f was derived to provide standardized effect size estimates. Pairwise comparisons were conducted with appropriate corrections for multiple testing.

For participants with complete methylation data, stress-induced changes in reproduction coefficients were calculated as Post-stress minus Pre-stress values and defined as ‘Stress change’. Gene × CIR Group interactions were examined using linear regression models: Stress change ∼ Gene × CIR Group + Sex + Age + BMI. This analysis was conducted separately for each task type and gene methylation (*COMT, DRD2, SLC6A3, TH),* testing whether the relationship between gene methylation levels and stress-induced performance changes differed between HCR and LCR groups.

For methylation analyses, outliers were identified using the same IQR method as behavioral data. Sensitivity analyses were conducted by re-examining gene × environment interactions after removing participants with extreme methylation values to assess robustness of findings.

## 3. Results

### 3.1. The Effect of TSST on State Anxiety and Cortisol Response

Before examining the effects of stress on time and space reproduction, we first verified that the TSST successfully induced psychosocial stress in our sample. Descriptive statistics for cortisol concentrations at all six sampling points are provided in Supplementary Table 2, and the group-averaged response curve is displayed in Figure 1B. As intended, cortisol levels rose following stress exposure and reached a peak approximately 15 minutes after the TSST. Specifically, concentrations at t_+30_ were significantly higher than those obtained immediately prior to stress induction (t_0_) [Z = –4.446, p <0.001; see Figure 1B]. Self-reported anxiety scores demonstrated a similar pattern. Participants’ state anxiety scores increased significantly from pre-TSST (34.25 ± 7.55) to post-TSST (46.48 ± 10.58) [*t*(43) = –9.220, 95% CI (–14.90, –9.55), p < 0.001]. Together, the robust increases in both state anxiety and cortisol confirm the effectiveness of the TSST in eliciting acute psychosocial stress, consistent with its established validity. The mean CIR across participants was 0.34 ± 0.41, ranging from –0.55 to 1.28. When examined by sex, mean CIR values were 0.30 ± 0.44 for men and 0.37 ± 0.40 for women. Although women showed a slightly higher mean CIR, this difference was not statistically significant [t(42) = 0.567, 95% CI (–0.18, 0.33), p = 0.567].

### 3.2. Primary Mixed-Effects Analysis

#### 3.2.1. Main Effects

The mixed-effects model revealed a dominant main effect of task type (*F*(1, 4108) = 790.44, p<0.001, f = 0.44), indicating substantial differences between temporal and spatial reproduction tasks. A significant main effect of stress emerged (*F*(1, 4102) = 21.72, p < 0.001, f = 0.07), showing overall performance changes following stress exposure. CIR Group showed a significant main effect (*F*(1, 38) = 4.50, p = 0.040, f = 0.03), indicating differences between high and low cortisol responders.

#### 3.2.2. Two-Way Interactions

The stress × task type interaction was significant (*F*(1, 4102) = 21.75, p < 0.001, f = 0.07), revealing domain-specific stress effects. Temporal reproduction improved under stress (Pre: M = −0.200, Post: M = −0.148; Δ = 0.052, SE = 0.008, t = −6.56, p < 0.001), while spatial reproduction remained unchanged (Pre: M = −0.015, Post: M = −0.015; Δ = 0.0002, SE = 0.008, t = 0.002, p = 0.998).

We observed a significant stress × sex interaction (*F*(1, 4102) = 4.46, p = 0.035, f = 0.03), indicating that men showed greater stress-induced performance changes than women overall. Post-hoc decomposition revealed this effect was driven primarily by temporal reproduction, where both sexes improved under stress but men showed substantially greater enhancement (Δ = 0.070, SE = 0.012, t = −5.96, p < .0001) than women (Δ = 0.035, SE = 0.011, t = −3.21, p = .0014). For spatial reproduction, neither sex showed significant stress-induced change (Men: Δ = −0.006, p = .614; Women: Δ = 0.006, p = .595). While the sex difference appeared more robust for temporal tasks, the three-way interaction did not reach significance (*F*(1, 4102) = 1.17, p = 0.279) suggesting the effect was present across both tasks (Figures 1C and 1D).

Men and women also showed differential performance profiles across tasks, as indicated by a significant sex × task interaction (*F*(1, 4108) = 12.30, p < 0.001, f = 0.05), independent of stress condition. Similarly, the CIR group × task type interaction (*F*(1, 4108) = 10.73, p = 0.001, f = 0.05) showed that high and low cortisol responders had different baseline performance patterns across temporal versus spatial tasks.

Other interactions were not significant: stress × CIR group (*F*(1, 4102) = 2.25, p = 0.133), CIR group × sex (*F*(1, 38) = 0.00, p = 0.969), stress × CIR group × task type (*F*(1, 4102) = 0.22, p = 0.639), and the four-way interaction (*F*(1, 4102) = 0.25, p = 0.617).

### 3.3. Gene × Cortisol Interactions

Analysis of methylation data (n = 88 observations from 44 participants) revealed gene × CIR group interactions specifically for temporal reproduction (Table 3). *DRD2* showed a significant interaction with CIR group for temporal tasks (β = 0.131, SE = 0.058, *t* = 2.28, p = 0.029), indicating that the relationship between *DRD2* methylation levels and stress-induced changes differed between HCR and LCR groups (Figure 2A and 2B).

**Figure 2.**
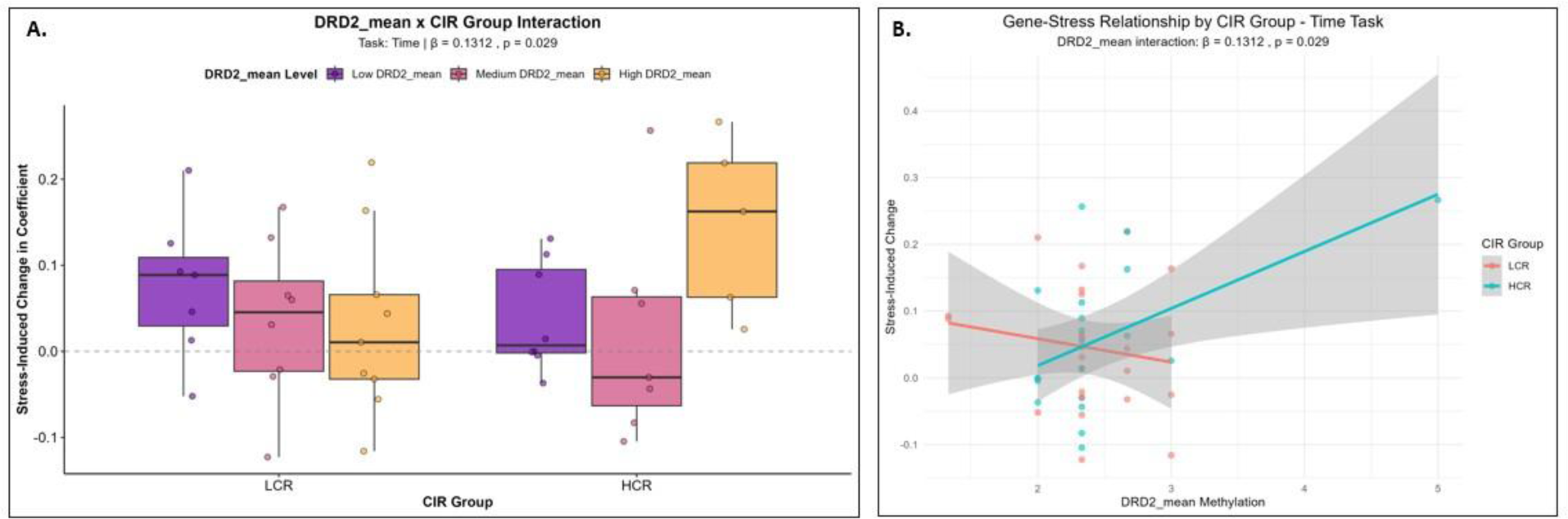
**A.** Stress-induced performance change and *DRD2* methylation levels by stress response groups (tertiles are created for visualisation) **B.** Stress-induced performance change by *DRD2* methylation levels for high and low cortisol responders.

**Table 3.**
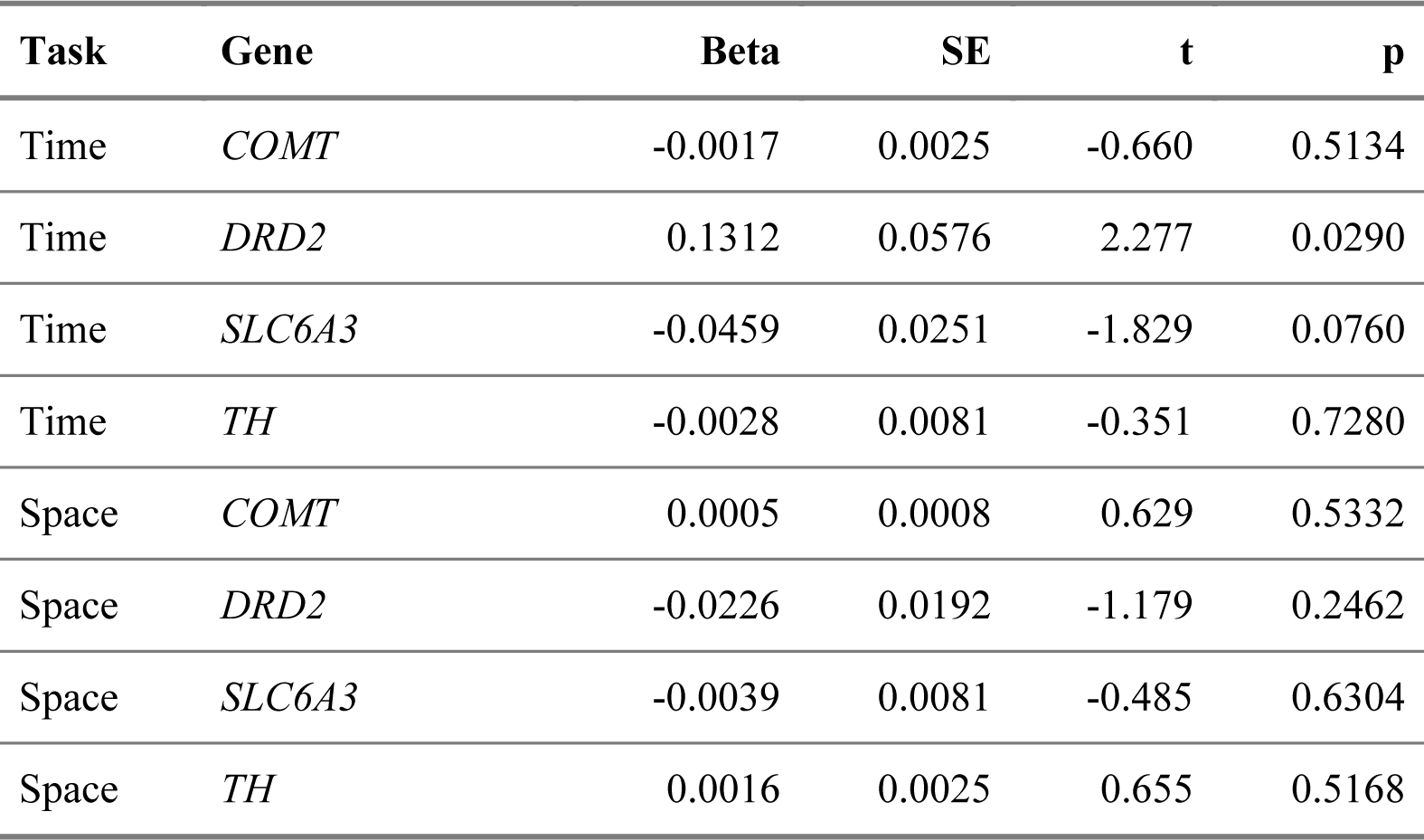
Gene by CIR Group interaction effects on stress-induced changes in reproductions.

To assess the robustness of these findings, we conducted sensitivity analysis by identifying methylation outliers using the same IQR criteria applied to behavioral data. *DRD2* showed 3 outliers (6.8% of participants). When outliers were excluded, effect size was attenuated (*DRD2*: β = 0.239, p = 0.067), indicating that this interaction requires replication in larger samples to establish robustness. No interactions emerged for spatial reproduction tasks (all p > 0.24).

### Integrated Findings

The results reveal a complex pattern where: (1) men show greater stress-induced improvement than women across both task types; (2) temporal processing specifically benefits from stress while spatial processing remains unaffected; (3) epigenetic variation in *DRD2* showed initial evidence of moderating stress-timing relationships, though sensitivity analysis indicated these effects require replication in larger samples to establish robustness; and (4) cortisol response patterns alone do not predict stress effects, but the potential role of dopaminergic gene methylation in temporal processing under stress requires further investigation.

## 4. Discussion

The present study investigated how acute psychosocial stress alters temporal and spatial reproduction and whether dopaminergic gene methylation moderates these effects. The TSST successfully elevated both subjective state anxiety and salivary cortisol, confirming effective stress induction. Behaviorally, participants showed the expected baseline underestimation of supra-second intervals and greater variability in time compared with space judgments, consistent with Vierordt’s law and prior findings that temporal reproduction is less precise than spatial reproduction (Droit-Volet et al., 2008; Murai and Yotsumoto, 2016; Robinson and Wiener, 2021). Critically, stress exposure led to improved temporal reproduction accuracy by reducing underestimation, whereas spatial reproduction remained unchanged. Exploratory analyses revealed that inter-individual differences in *DRD2* (D2 receptor) methylation moderated temporal responses to stress, highlighting a role for dopamine-related epigenetic regulation.

### 4.1. Stress and time perception

Our finding that stress reduces the tendency to underestimate time, thereby improving accuracy, builds on previous research into stress and interval timing. Much of the existing literature has emphasized time dilation, the subjective lengthening of intervals when individuals face threatening or emotionally negative stimuli. For example, fearful or unpleasant cues reliably produce overestimation of duration (Bar-Haim et al., 2010; Droit-Volet et al., 2004). These results support models in which rapid autonomic arousal accelerates an internal pacemaker, thereby lengthening perceived duration (Mella et al., 2011).

However, relatively few studies have examined how post-stress states affect timing. Van Hedger et al. (2017) reported that social stress produced dilation in verbal estimation tasks, but effects were not explained by autonomic reactivity. Yao et al. (2016) found no overall effect of social stress but observed reduced temporal sensitivity among stronger cortisol responders; although limited by an all-male sample and the absence of a control condition. Similarly, Kale et al. (2019) found no significant effect of fearful facial expressions on time discrimination sensitivity, suggesting that the presence of threatening stimuli alone may not be sufficient to alter temporal processing when attention is divided. Our findings diverge from these reports: we observed improved accuracy in supra-second reproduction following stress. Several factors may account for this discrepancy. First, we tested performance during the expected cortisol peak, targeting the slower-acting HPA axis rather than immediate SAM activity. Second, our reproduction paradigm required not only perceptual encoding but also retention and motor reproduction, processes that are heavily dependent on corticostriatal circuits. Finally, we included a matched spatial control, strengthening the interpretation that effects were specific to temporal processes rather than general arousal or performance shifts.

The domain specificity of stress effects aligns with longstanding evidence that interval timing depends on dopaminergic signalling. Lesion, neuroimaging, and patient studies consistently implicate basal ganglia circuits in temporal processes (Coull et al., 2011; Merchant et al., 2013). Parkinson’s disease, marked by dopamine loss in the nigrostriatal pathway, is characterized by systematic impairments in interval timing (Merchant et al., 2008; Smith et al., 2007). Pharmacological studies show that dopaminergic agonists and antagonists systematically bias reproduced durations (Rammsayer 1989, 1993, 1997; Rammsayer and Vogel 1992). D2 receptor activity plays a particularly central role in regulating interval timing (Rammsayer, 2009), and optogenetic manipulations of dopamine neurons provide causal evidence that direct modulation of dopaminergic firing can immediately and transitively bias time perception (Soares et al., 2016).

Within this framework, our finding of dilated reproduction following stress is compatible with models in which stress hormones influence dopaminergic signalling in corticostriatal circuits (Belujon and Grace, 2015). GRs are highly expressed in striatum, hippocampus, and prefrontal cortex (Arnsten, 2009; Piazza and Le Moal, 1998), providing potential routes for cortisol-dopamine interactions, though the specific mechanisms linking HPA activation to temporal processing remain to be determined.

### 4.2. Epigenetic moderation

Another novel contribution of this study is the preliminary evidence that epigenetic variation in dopamine-related genes may shape how stress influences timing. Initial analysis revealed that *DRD2* methylation significantly interacted with CIR group, such that higher methylation was associated with greater stress-induced change in temporal reproduction among high cortisol responders. However, sensitivity analysis revealed that this interaction was primarily driven by participants with extreme methylation values, as effects were substantially weakened when statistical outliers were removed. This finding raises questions about interpreting epigenetic variability in behavioral research, as participants with extreme methylation values may represent meaningful biological variation rather than statistical artifacts. However, the small sample size limits our ability to distinguish between these interpretations, and replication with larger samples is essential.

Mechanistically, higher *DRD2* methylation likely reduce postsynaptic D2 receptor expression, thereby altering the sensitivity of striatal circuits to dopaminergic modulation under stress. This interpretation is consistent with pharmacological evidence that D2 receptor antagonism disrupts interval timing, while D2 stimulation enhances it (Rammsayer, 2009, 1997, 1993). The emergence of this effect most clearly in high cortisol responders suggests that stress physiology and dopaminergic receptor regulation interact in ways that may be particularly pronounced in individuals with specific methylation profiles. By contrast, *COMT, SLC6A3,* and *TH* methylation did not moderate temporal performance. This may reflect their broader, less direct roles in dopamine metabolism (synthesis, reuptake, and degradation) compared to the receptor-mediated signaling that appears critical for interval timing under stress.

Importantly, while promoter methylation is often associated with reduced gene expression (Moore et al., 2013), these relationships are CpG- and tissue-specific. Our saliva-based measures serve as peripheral proxies of central regulation. Recent evidence shows strong brain-saliva methylation correlations for dopaminergic genes (r>0.85; (Braun et al., 2019), supporting the validity of this approach. Though these peripheral measures do not directly reflect synaptic dopamine signaling, the moderation pattern we observe aligns with pharmacological and animal studies demonstrating the critical role of D2 receptor function in interval timing.

The present findings provide preliminary evidence that epigenetic variation of dopamine-related genes may moderate stress-induced shifts in human time perception, though the sensitivity of these effects to extreme methylation values indicates that larger samples are needed to establish robustness.

### 4.3. Sex differences and variability

We observed a significant stress × sex interaction, with men showing greater stress-induced performance changes than women overall. For temporal reproduction, both sexes showed improved accuracy under stress, but men demonstrated substantially greater enhancement (Δ = 0.070, p < .001) than women (Δ = 0.035, p = .001). For spatial reproduction, neither sex showed significant stress-induced changes (both p > .59). Although this sex difference was most evident in temporal tasks, the non-significant three-way interaction (p = .279) suggests that the magnitude of the sex difference did not reliably vary between temporal and spatial domains.

Importantly, this sex difference cannot be attributed to differential HPA reactivity, as cortisol reactivity did not differ between sexes in our sample. The mechanisms underlying these sex-related differences in stress-cognition interactions remain unclear and warrant further investigation, ideally through direct assessment of sex hormone levels and their interplay with stress-induced cognitive changes.

### 4.4. Limitations and Future Directions

Several limitations should be acknowledged. The sample size provided limited power for detecting epigenetic interactions, as evidenced by the sensitivity of methylation findings to individual participants with extreme values. Whether these participants represent statistical outliers or meaningful biological variation within the broader population remains unclear and highlights the fundamental challenge of studying epigenetics-behavior relationships in smaller samples.

The temporal paradigm was restricted to supra-second reproduction, limiting generalizability to other timing domains. We measured cortisol as our primary stress biomarker but lacked autonomic indices, preventing us from dissociating HPA vs. SAM contributions to observed effects. Claims about dopaminergic mechanisms remain inferential, as we did not directly assess neurotransmitter function. Saliva-based DNA methylation provides only peripheral estimates of central nervous system regulation.

Future research should prioritize larger, more diverse samples capable of representing broader spectrum of epigenetic variation. Multi-task approaches and integration of autonomic measures with cortisol sampling would provide a more complete picture of stress-timing relationships. Longitudinal designs tracking stress exposure and methylation patterns over time would address questions about adaptation and plasticity in these systems.

### 4.5. Conclusion

This study demonstrates that acute psychosocial stress selectively enhances temporal reproduction accuracy while leaving spatial processing unchanged at the group level. However, this domain-specific stress effect was moderated by sex in a domain-general manner: men showed stress-induced improvement across both temporal and spatial tasks, whereas women showed minimal change in either domain. Importantly, these sex differences could not be attributed to differential cortisol reactivity, as HPA responses did not differ between sexes.

Initial analyses suggested that *DRD2* methylation moderates stress effects on timing, though sensitivity analyses revealed these interactions were driven by participants with extreme methylation values. Whether these individuals represent meaningful biological variation or statistical artifacts cannot be determined in the current sample size, highlighting a key challenge for behavioral epigenetics research.

These results demonstrate that stress-cognition relationships involve multiple levels of specificity: stress produced domain-specific enhancement of temporal accuracy overall, yet individual differences in stress responsivity (related to sex) manifested domain-generally across both temporal and spatial performance. Preliminary evidence suggests that epigenetic variation in dopamine-related genes may further modulate these effects, though small sample size limits firm conclusions. This study provides a framework for behavioral epigenetics research and demonstrates that understanding stress-cognition relationships requires acknowledging individual biological variation in stress responsivity.

## Author Contributions

**G.G.A.** contributed to Conceptualization (study hypothesis and experimental design), Methodology (molecular protocols and behavioral framework), Investigation (wet lab experiments including DNA extraction, pyrosequencing, and cortisol ELISA), Data Curation (cortisol and methylation data), Formal Analysis (wet lab metrics and group comparisons), Resources, and Writing-Review&Editing. **G.V.** contributed to Methodology (behavioral protocols), Software (task and statistical analysis code), Investigation (conducting the TSST, data and sample collection), Data Curation (behavioral), Formal Analysis (multimodal integrations, mixed-effects models, sensitivity analyses), Visualization, Writing-Original Draft, and Writing-Review&Editing. **F.Ç.** contributed to Investigation (sample processing, wet lab). **N.V.** contributed to Investigation (pyrosequencing) and Writing-Review&Editing. **M.Ç.** contributed to Conceptualization, Supervision, Project Administration, Funding Acquisition, and Writing-Review&Editing.

## Data Availability Statement

*The behavioral and wetlab datasets, and analysis scripts supporting the findings of this study have been deposited in the Open Science Framework repository and can be accessed at* https://osf.io/s2bgm/ *[DOI: 10.17605/OSF.IO/S2BGM]*.

## Acknowledgements

This work was supported by the Scientific and Technological Research Council of Turkey (TUBITAK, Project No: 219K098). We would like to thank to İrem Öykü Kahraman and Ayşe Gökçe Erman for helping sample collection, also to Can Karahan and Şeyma Gül Durgut for technical support.

