## Supplementary for "Interval Timing Shows Selective Enhancement Under Psychosocial Stress: A Cortisol-Mediated Dissociation From Spatial Processing"

**Psychometric Measures**

**TSST Background Questionnaire and Sociodemographic Evaluation**

In order to confirm the restrictions required for the TSST application (coffee consumption, smoking, alcohol intake, and drug use) a detailed questionnaire suggested by Linares et al (2020) for making the TSST methodology more systematic and detailed in reporting was used when participants arrived to the experiment (Narvaez Linares et al., 2020). TSST background questionnaire requires the participants to report on their physical exercise, alcohol, caffeinated beverages, tobacco products, recreational substances and food consumption, and tooth brushing 24 hours before the experiment. These questions assess the extent to which the participant adheres to the pre-experimental guidelines. In addition to these restrictions, it allows experimenters to screen menstrual cycle of the women participants, and document the participant's sociodemographic characteristics and medical status (e.g. education, height, weight, chronic diseases). Due to the experiments took place during COVID-19 outbreak months, we also added COVID-related questions to this questionnaire such as COVID-19 vaccination status and history of COVID-19 infection.

**Handedness Scale (HS)**

This questionnaire, developed by Chapman and Chapman (1987), is designed to ascertain which hand people prefer to use in their daily lives for a range of activities, including using a hammer, brushing teeth and using scissors (Chapman and Chapman, 1987). The questionnaire comprises 13 items, with 1 point awarded for a right hand response, 3 points for a left hand response, and 2 points for both hand responses. A score between 13 (fully right-handed) and 39 (fully left-handed) is therefore obtained. The validity and reliability of the Turkish adaptation were conducted by Nalçacı et al. (2002) (Nalçacı et al., 2002).

**Spilberger State-Trait Anxiety Inventory (STAI)**

The scale was developed by Spielberger et al. (1970) as a self-report instrument for the assessment of state and trait anxiety. The scale comprises two distinct sections, each comprising 20 items, with a scoring range of 0-4. A high score is indicative of elevated anxiety levels (Spielberger et al., 1970). Oner and Le Compte (1985) conducted a validation and reliability study for the Turkish population (Oner and Le Compte, 1998).

**Trail Making Test (TMT)**

The Trail Making Test, initially developed by Reitan et al. (1955), is a neuropsychological assessment tool commonly used to evaluate executive functions such as working memory, complex attention and planning (Reitan, 1958). The test comprises A and B forms. The A section of the test assesses visual scanning-based processing speed. The B section assesses the ability to change the set-up between stimulus sets and to follow the succession. The test is scored based on the number of correct responses and the time taken to complete the forms. Türkeş et al. (2015) conducted a Turkish validity and reliability study of the test. In the current study B/A time was calculated and used (Türkeş et al., 2015).

**Perceived Stress Scale (PSS)**

The Perceived Stress Scale was developed by Cohen et al. (1983) to evaluate the extent to which individuals perceive specific life situations as stressful and how stressful people perceive various situations in their lives (Cohen et al., 1983). This test, of which validity and reliability studies in Turkish were conducted by by Eskin et al. (2013), comprises 14 items, each scored between 0 and 4, with 7 items scored in reverse order. The total score ranges from 0 to 56, with higher scores indicating greater stress levels (Eskin et al., 2013).

**Liebovitz Social Anxiety Scale (LSAS)**

The Liebowitz (1987) scale is a tool used to assess fear and avoidance during social interaction and performance in individuals with social anxiety disorder. Two sub-scores, anxiety and avoidance, can be calculated with this scale, which is also suitable for use with the general population. It consists of a total of 24 items and higher scores indicate greater anxiety and avoidance (Liebowitz, 1987). The Turkish version of the scale was adapted by Soykan et al. (2003) (Soykan et al., 2003).

**Adult Attention Deficit Hyperactivity Disorder Self-Report Scale (ASRS)**

The Adult ADHD Self-Report Scale was developed in collaboration with the World Health Organization and the Adult ADHD Study Group. It is designed to assess attention deficit hyperactivity disorder (ADHD) in adults through self-report, making it suitable for epidemiological studies (Kessler et al., 2005). The 18-item scale comprises two subscales, each comprising nine items designed to assess symptoms of attention deficit and hyperactivity/impulsivity. Each item enquires about the frequency of symptoms experienced over the previous six months. The five-point Likert-type scale allows for a minimum score of 0 and a maximum score of 72 to be obtained. The Turkish validity study of the scale was conducted by Doğan et al. (2009) (Doğan et al., 2009).

**Beck Depression Inventory (BDI)**

The Beck Depression Inventory (BDI), developed by Beck et al. (1988), is a 21-item scale designed to assess the symptoms of depression. Each item is scored on a scale of 0-3, with a total score of 0-63 achievable on the scale. A higher total score indicates a greater severity of depressive symptoms (Beck et al., 1988). Hisli et al. (1989) adapted the scale into Turkish (Hisli, 1989).

**TRIER Social Stress Test**

The TSST is a highly ecologically valid acute stress protocol that is considered the gold standard for experimentally studying the stress response in humans. It was developed by Clemens Kirschbaum. In this test, individuals are forced to speak in front of an unresponsive jury and are stressed by social evaluation, unexpected events and public speaking factors. The TSST has been used as a stressor in many studies in the literature and has been shown to activate the HPA axis (Kirschbaum et al., 1993; Yamanaka et al., 2019).

The participant, who had not been given any prior information or instructions about the TSST, was immediately escorted to the TSST room following the conclusion of the initial phase of the time perception paradigm. Candidates were asked to prepare a five-minute speech to convince the jury of their suitability for the job in a scenario where they applied for a position in their desired field. The participant was informed that the speeches would be delivered in front of the microphone, recorded on video, and subsequently analyzed by a competent team for behavioral and tone of voice aspects. However, nothing was actually recorded. The participants were then permitted to prepare alone in the TSST room for a period of five minutes. At the end of this period, after the jury had taken their seats, the test began. The participants were requested to begin their speech. During the course of the test, the members of the jury did not display any gestures or mimics that could be interpreted as positive or negative feedback. Furthermore, they did not respond to the questions of the participants. If they finished their presentations before the five-minute limit, the jury issued a clear warning and asked them to continue. Once the speaking component was concluded, the jury presented the participant with an unexpected mental arithmetic challenge, which they were required to complete for a period of five minutes. In this task, the participant was required to count backwards by 13, beginning with the number 1022. Upon each instance of an incorrect response, the participant was given a warning, "You made a mistake. Please begin counting from the beginning," and was asked to restart. The arithmetic task was also concluded at the end of five minutes.

**Saliva Sampling**

Saliva samples were collected from the participants at six different time points (t_-30_, t_0_, t_+15_, t_+30_, t_+45_ and t_+60_) in order to determine the cortisol response. To this end, Salivette® Cortisol tubes (Sarstedt, 51.1534.500), comprising a cylindrical cotton-like synthetic material called a swab, were utilized. At each time point, the Salivette® swab was placed inside the participant's cheek and saliva was absorbed by the swab for two minutes. The swabs were then transferred to their own tubes and centrifuged at 3000 rpm for 15 minutes. The supernatant was aliquated into three sterile microcentrifuge tubes and stored at -80°C until the ELISA was performed.

| **Supplementary Table 1.** Methylation specific PCR primers for *DRD2, COMT, TH and SLC6A3* genes. | | | | |
| --- | --- | --- | --- | --- |
| ***Gene Symbol*** | | **Gene Name** | **NCBI Gene ID #** | **Primer Information** |
| ***DRD2*** | Dopamine receptor D2 | | 1813 | The primers were selected from the PyroMark CpG assay, GeneGlobe catalogue (Qiagen, Hs_DRD2_01_PM, PM00047411). *The manufacturer does not make the primer information available.* |
| ***COMT*** | catechol-O-methyltransferase | | 1312 | Forward (5’-3’) TGG GGA GAG TTG TTA GTT GTT ATA AGT ATT  Reverse (5’-3’) TCA AAA CAA TTC AAA CCT CAT CCT ATA A  Sequencing (5’-3’) ATG GTG GGG GAT TAT  *The primer sequences were sourced from Gao et.al. (2017)(Gao et al., 2017).* |
| ***TH*** | Tyrosine hydroxylase | | 7054 | Primers were selected from the PyroMark CpG assay, GeneGlobe catalogue (Hs_TH_01_PM, PM00155750). *The manufacturer does not make the primer information available.* |
| ***SLC6A3*** | Solute carrier family 6 member 3, Dopamine transporter | | 6531 | Primers were selected from the PyroMark CpG assay, GeneGlobe catalogue (Hs_SLC6A3_01_PM, PM00022064). *The manufacturer does not make the primer information available.* |

| **Supplementary Table 2**. Descriptive statistics for cortisol values of the study participants (n=44) corresponding to six different time points. | | |
| --- | --- | --- |
|  | **Cortisol level (nmol/L)** | |
| **Time point (minutes)** | **Mean (SD)** | **Median (Minimum-Maximum)** |
| t_-30_ | 18.63 (14.58) | 15.30 (4.25-86.80) |
| t_0_ | 20.92 (14.46) | 16.00 (5.95-62.46) |
| t_+15_ | 22.87 (14.14) | 19.22 (5.50-56.63) |
| t_+30_ | 30.96 (17.01) | 30.11 (6.56-75.25) |
| t_+45_ | 25.47 (13.87) | 22.78 (4.00-59.49) |
| t_+60_ | 22.04 (11.27) | 19.07 (3.78-47.89) |
| CIR | 0.34 (0.41) | 0.26 (-0.55-1.28) |
| CIR: Cortisol increase rate, SD: Standard deviation | | |

| **Supplementary Table 3.** Descriptive Statistics and Intergroup Comparison Results for Demographic Data and Psychometric Measurements of LCR and HCR Groups | | | | | | |
| --- | --- | --- | --- | --- | --- | --- |
|  | **LCR (N=24)** | | **HCR (N=20)** | | **Intergroup comparisons** | |
| **Demographics and Psychometric Variables** | Mean  (SD) | Median  (Min-Max) | Mean  (SD) | Median  (Min-Max) |  |  |
| Age (years) | 22,29  (3,42) | 21,00  (19,00-32,00) | 21,55  (2,52) | 21,50  (18,00-28,00) | U=220,00; P=0,663^a^ | |
| Education (years) | 16,67  (3,23) | 16,00  (14,00-26,00) | 16,15  (1,76) | 16,00  (13,00-20,00) | | U=229,00; P=0,792^a^ |
| BMI | 22,34  (2,67) | 22,50  (18,10-28,10) | 21,74  (2,44) | 21,45  (17,20-26,00) | | t(42)=0,769; P=0,418^b^ |
| Trail Making Test B/A | 2,25  (0,90) | 2,03  (1,27-5,32) | 2,18  (0,52) | 2,03  (1,19-2,28) | | U=230,50; P=0,823^a^ |
| STAI-*Trait Anxiety* | 43,42  (11,27) | 44,50  (22,00-62,00) | 42,25  (9,78) | 39,00  (27,00-59,00) | | t(42)=0,363; P=0,719^b^ |
| STAI-*State Anxiety* _pre-TSST_ | 34,04  (8,46) | 32,50  (20,00-52,00) | 34,50  (6,52) | 34,00  (22,00-47,00) | | t(42)=-0,198; P=0,844^¶^ |
| STAI- *State Anxiety* _post-TSST_ | 43,46  (10,01) | 42,50  (20,00-58,00) | 50,10  (10,26) | 51,00  (32,00-67,00) | | **t(42)=-2,161; P=0,036**^b^ ***** |
| Liebovitz Social Axiety Scale- *Anxiety* | 23,96  (13,28) | 22,50  (6,00-50,00) | 26,55  (9,82) | 29,50  (13,00-39,00) | | U=202,00; P=0,370 ^a^ |
| Liebovitz Social Axiety Scale - *Avoidance* | 18,58  (10,74) | 16,50  (1,00-38,00) | 22,35  (9,53) | 20,50  (9,00-46,00) | | t(42)=-1,218; P=0,230^b^ |
| Liebovitz Social Axiety Scale - *Total* | 42,54  (22,83) | 38,50  (7,00-88,00) | 48,90  (18,30) | 48,00  (22,00-84,00) | | t(42)=-1,005; P=0,321^b^ |
| Beck Depression Inventory | 12,79  (9,44) | 12,00  (0,00-35,00) | 10,65  (6,95) | 8,50  (2,00-25,00) | | U=206,00; P=0,422 ^a^ |
| Percieved Stress Scale | 25,08  (8,15) | 24,50  (4,00-39,00) | 26,50  (7,28) | 26,50  (16,00-40,00) | | t(42)=-0,602; P=0,550^b^ |
| Adult Attention Deficit Hyperactivity Disorder Self-Report Scale | 33,00  (8,58) | 33,50  (10,00-50,00) | 32,50  (9,27) | 33,50  (15,00-55,00) | | t(42)=0,186; P=0,854^b^ |
| BMI: body mass index, HCR: High cortisol responders, LCR: Low cortisol responders, pre-TSST: before the TRIER social stress test, post-TSST: after the TRIER social stress test, SD: standard deviation.  ^a^ Mann-Whitney U Test, ^b^ Independent-Samples T-test | | | | | | |

| **Supplementary Table 4.** Descriptive statistics of % methylation levels and number of CpG sites analysed in *DRD2*, *COMT*, *TH*, and *SLC6A3* genes. | | | |
| --- | --- | --- | --- |
| **Gene of Interest** | **Number of CpG sites anlysed** | **Avarage Methylation Level (%)** | |
|  |  | **Mean (SD)** | **Median (Min-Max)** |
| ***DRD2* (N=44)** | 3 | 2.42 (0.54) | 2.33 (1.33-5.00) |
| ***COMT* (N=44)** | 3 | 64.05 (15.67) | 69.33 (4.33-7.33) |
| ***TH* (N=43)** | 5 | 80.02 (4.52) | 80.60 (78.00-93.40) |
| ***SLC6A3* (N=42)** | 6 | 8.40 (1.50) | 8.83 (2.83-10.50) |
| SD: Standard deviation | | | |

| **Supplementary Table 5.** Descriptive statistics of demographic and psychometric measures (N=44). | | |
| --- | --- | --- |
| **Demographic and Psychometric Variables** | **Mean (SD)** | **Median (Minimum-Maximum)** |
| Age (years) | 21.96 (3.03) | 21.00 (18.00-32.00) |
| Education Duration (years) | 16.43 (2.65) | 16.00 (13.00-26.00) |
| BMI | 22.07 (2.56) | 21.65 (17.20-28.10) |
| Trail Making Test B/A | 2.22 (0.74) | 2.03 (1.19 – 5.32) |
| STAI-*Trait Anxiety* | 42.89 (10.51) | 40.00 (22.00-62.00) |
| STAI-*State Anxiety* _pre-TSST_ | 34.25 (7.55) | 34.00 (20.00-52.00) |
| STAI- *State Anxiety* _post-TSST_ | 46.48 (10.58) | 46.00 (20.00-67.00) |
| Liebovitz Social Axiety Scale- *Anxiety* | 25.14 (11.77) | 26.50 (6.00-50.00) |
| Liebovitz Social Axiety Scale - *Avoidance* | 20.30 (10.27) | 19.00 (1.00-46.00) |
| Liebovitz Social Axiety Scale - *Total* | 45.43 (20.91) | 44.00 (7.00-88.00) |
| Beck Depression Inventory | 11.82 (8.863) | 8.38 (0.00-35.00) |
| Percieved Stress Scale | 25.73 (7.71) | 25.00 (4.00-40.00) |
| Adult Attention Deficit Hyperactivity Disorder Self-Report Scale | 32.77 (8.80) | 33.50 (10.00-55.00) |
| BMI: body mass index, pre-TSST: before the TRIER social stress test, post-TSST: after the TRIER social stress test, SD: standard deviation | | |

| **Supplementary Table 6.** Correlation analysis between DNA methylation of dopamine related genes and CIR | | |
| --- | --- | --- |
| Correlations  Spearman rho; P value | | CIR |
| % methylation | *DRD2 (N=44)* | -0.053 ;0.738 |
|  | *COMT (N=44)* | 0.039; 0.804 |
|  | *SLC6A3 (N=42)* | **0.339; 0.030 *** |
|  | *TH (N=43)* | 0.247; 0.115 |
| Control variable: Gender , CIR: Cortisol incease rate | | |
